# Looking at the map – or navigating in a virtual city interaction of visuospatial display and spatial strategies in VR

**DOI:** 10.1101/405233

**Authors:** Zsolt Gyozo Török, Ágoston Török

## Abstract

To study geo-visualization processes a Cognitive Cartography Lab was established at Eötvös University, and the “Virtual Tourist” experiment was designed for the better understanding of actual map use during navigation. In this paper we present some preliminary results of the experiment. We explored the use of a static, north-oriented city map during navigation in an interactive, 3D town. Participants explored the virtual environment or followed verbal instructions before they completed spatial tasks. Their spatial behavior, verbal reactions were recorded, and also eye tracking data from 64 participants was collected. The experiment was designed by a multidisciplinary research group, including students of Eötvös Loránd University.

## I. INTRODUCTION

Humans have been using visuospatial displays, external cognitive tools since millennia [1]. Since ancient times we use maps as cognitive tools to visualize complex spaces, but only in the last centuries are maps routinely used to help everyday wayfinding and navigation in geographic space. Despite the long history of cartography, little is known about why and how visuospatial displays, especially maps help cognitive processes, including spatial learning. Starting with the first maps to modern days GPS – cognitive tools help us in orienting, wayfinding and navigating in geographic space, at the same time, these cognitive instruments reflect the plasticity and changing functionality of the human brain.

Cognitive infocommunication is an emerging interdisciplinary field integrating human cognitive and ICT technologies into new synergies [2]. As recent research demonstrates – despite the availability of global satellite navigational systems (GNNS) – the development and practice of human spatial abilities by traditional navigational tasks, that is reading a map, have key role in efficient human-computer interaction with all kinds of dynamic and adaptive information and data visualization [3]. Moreover, studies on spatial learning suggest that people who successfully build cognitive maps, or labeled graphs in their memory are better with STEM subjects in schools. so spatial thinking is one of the most important fields for the development of education in modern human society [4].

Siegel and White proposed that human spatial knowledge acquisition follows a hierarchical structure [5]. First, when we encounter with a novel environment, we acquire *landmark knowledge* that contains distinctive elements of the environment or scenes stored in memory [6]. In literature landmarks sometimes are categorized as local or global ones [7]. Former one can be observed only from a close distance, like a shop or a building. To local landmarks intermediate goals are related, such as turning to a specified direction, thus they are linked to route navigation. In contrast, global landmarks refer to a well distinguishable point of reference from a broader viewpoint, therefore they can suggest a geocentric (geographic) framework for navigation [8].

The next level in the hierarchy is *route knowledge*. It consists landmarks, places and sequential turns or directions attached to them and route connections between them. Route knowledge enables us to follow an already known path from one point to another destination, containing sequential elements of directions, like a turn to right then a turn to the left at the given landmark [9].

Finally, if we gain an integrated knowledge about the environment by time, by experience and/or by other external aids such as maps, we can acquire the highest or most advanced level of spatial representation, *survey knowledge* [10]. This internal representation of a large-scale, geographic environment was called ‘*cognitive map*’, a function of the sophisticated memory structure created the hippocampus and parahippocampal areas in the human brain [11] [12]. Survey knowledge helps more in effective navigation and route planning [13] [14], in contrast to route knowledge, because it provides an external reference frame which leads to an overview of the spatial layout, and this enables us to take shortcuts or to navigate in unfamiliar routes.

The high importance of *reference frame* in human navigation is underlined by recent neuropsychological research, showing their dependence of viewpoint [15]. Furthermore, this integrated spatial representation provides metric information, such as the relationships between different locations and landmarks [16]. The type of spatial knowledge which can be acquired in a novel environment depends on the goals of learning, or on the effort taken into consideration or on the structure of the environment [17].

Furthermore, as mentioned above, humans tend to use different graphic representations to facilitate their navigation spatial knowledge can be developed by using overview methods, such as maps.

However, not every type of navigation assistance contributes to the same aspects of spatial knowledge. We can make a distinction between static spatial representations and dynamic ones, the former category can refer to the classic paper-based, north-oriented maps with allocentric (or geographic) reference frame, while the latter one here refers to novel navigational systems such as GPS-based and mobile apps. These latter systems can track and visualize the spatial location of the user and habitually offer a forward-up (egocentric) orientation, which means that they rotate themselves according to our headings [18].

## II. THE VIRTUAL TOURIST EXPERIMENT

### A. Participants

Sixty-four participants took part in the experiment, data from nine participants was excluded due to technical errors in recording. The analysis was therefore run on data from fifty-five participants (30 female). Their age ranged from 18 to 25 (M = 20.49, SD = 2.00). Their vision was normal or corrected to normal.

All participants were first year bachelor students from the Eötvös Loránd University, either from the Department of Cartography and Geoinformatics, from the Department of Geography, or from the Faculty of Education and Psychology. The experiment was approved by the research ethical board of the Eötvös University. Beyond the opportunity to gain firsthand experience from cognitive research the participants received bonus points on their exam performance for their activity. Prior to the experiment, participants were given explanation and they gave a written informed consent.

### B. Apparatus and stimuli

The Virtual Tourist VR experiment was designed with the adoption of the concept gamification [18]. The city map was designed with a legend following Google map style (2018) because of its familiarity and popularity, using the CorelDRAW x8 graphic design software. The map was based on an old, instructional topographic map sheet of an imaginary region but was re-designed in the style Google maps. The printed map was made as a military educational map and used in the Socialist era (1959) [19]. This base map was used as a basis of our design, which was actually a compilation of map details of different regions in Hungary to make an ecologically valid representation of different types of terrain by the actual set of conventional signs. In the period when topographic maps were secret it could be used to train general map reading without restrictions. While a familiar environment, a typical Central European landscape and cityscape the representation of the fictional town ‘Szegvár’ was unknown to all test subjects.

We used a smaller central part (∼1×1 km) of the original map to build up a 3D environment in Unity. The streets were named after famous Hungarian scientists and artists, otherwise, the town consisted of three different building types: houses, shops, and unique buildings, “Museums” and the Cathedral standing on the squares (see layout on Figure 1).

**Figure 1.**
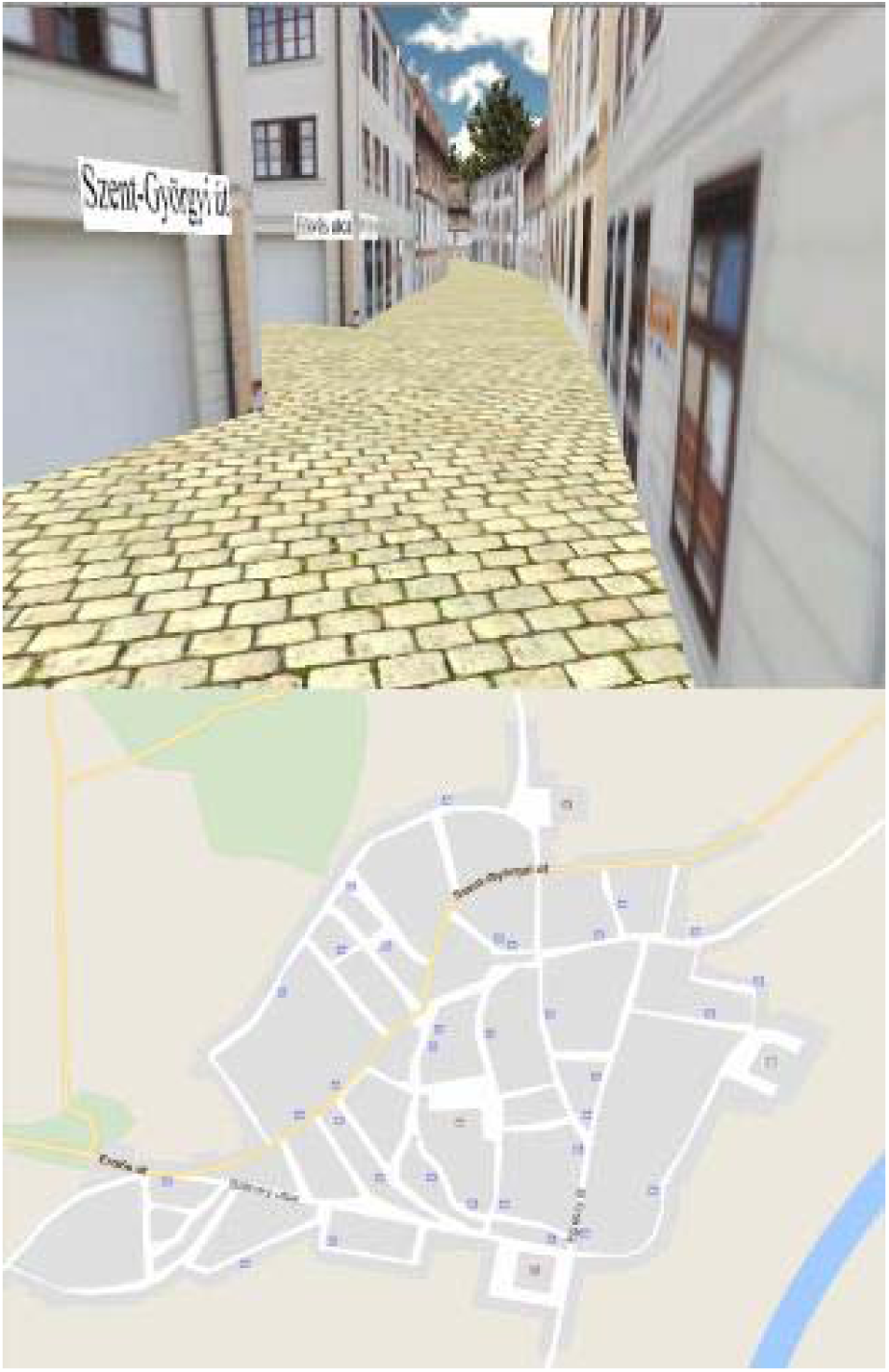
The layout of scree in the virtual environment: the interactive, realistic city-model in the upper half, the conventional cartographic representation in the lower part part of the upright format screen

The virtual reality game was programmed in Unity 4 (4.6.7f1) and it was presented on a 9:16 aspect ratio screen with a resolution of 1080×1920 pixels. The participants sat in front of the screen in a distance approximately 55 centimeters. For the navigation in the virtual environment they could use a keyboard placed in front of them. The game’s frame was a fictional visit as tourists to a town called ‘Szegvár’, where the participants could explore the city by themselves or with the help of a tourist guide (Exploration phase) and then they could buy some souvenirs (Test phase). To approximate actual map use in the field we considered the real situation of a real tourist at ground level, looking forward and navigating, making spatial decisions derived from the analysis of visual information (scene recognition, path optimization). As in real situations, the ‘head-up-is-forward’ position during navigation, we decided a layout that imitated the practical arrangement of the map as an object in the lower part of the visual space.

The town was shown in the upper half of the screen (egocentric reference frame), while in the bottom half of the screen a map was presented (allocentric reference frame) (Figure 3.) With constant walking speed, the city’s diameter was navigable in its major road in approximately five minutes.

**Figure 2.**
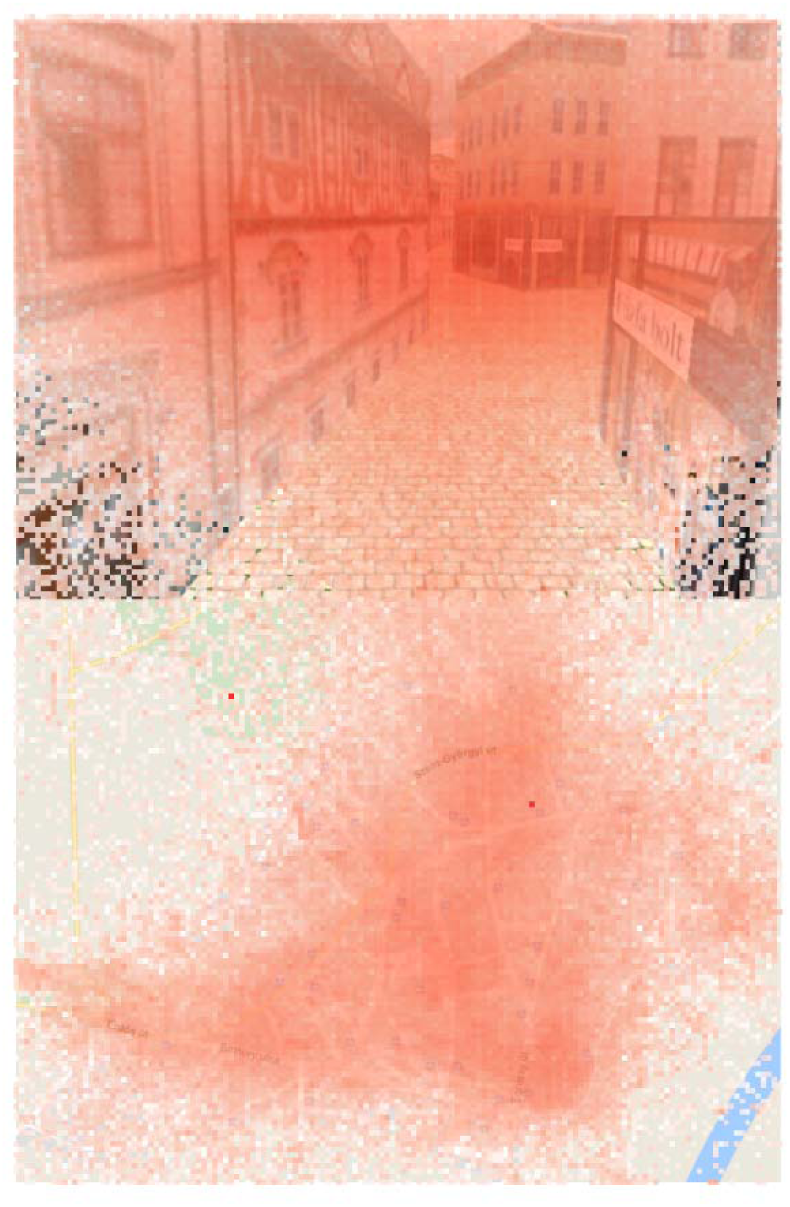
Heat map of fixations from all participant overlaid on the image of the screenshot from the VR experiment.

**Figure 3.**
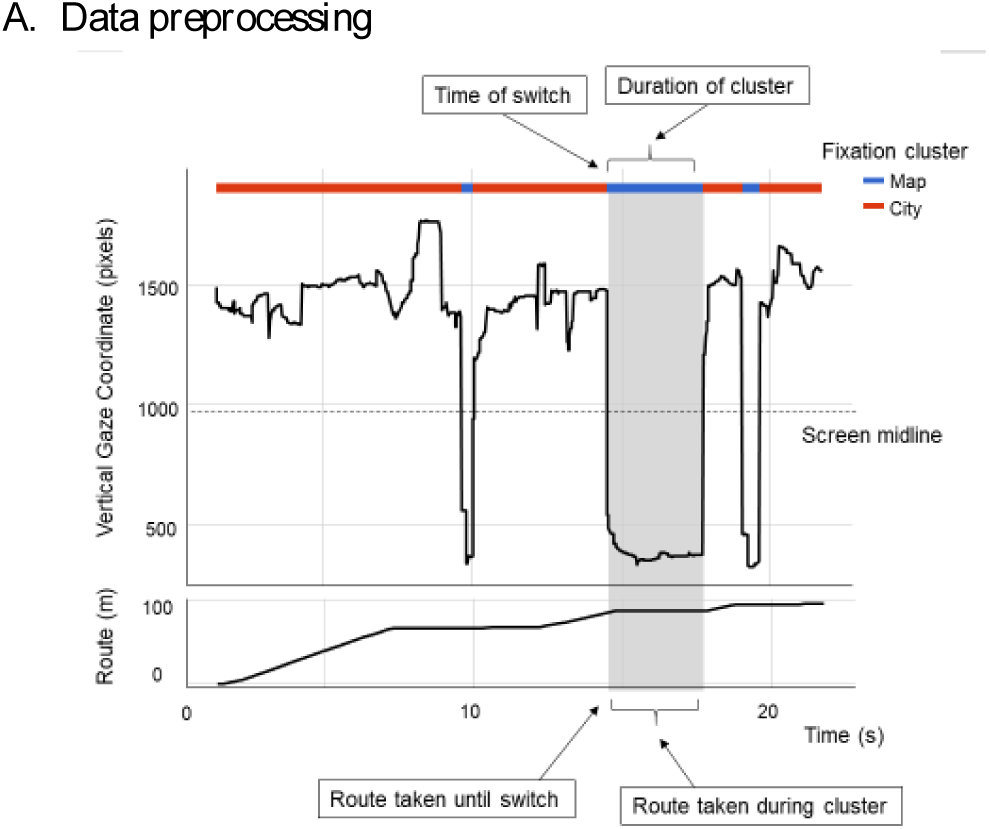
The plotting of the vertical coordinates of the gaze of a participant into a spatio-temporal graphic space reveals periods of walking when looking at the map

During the experiment we recorded gaze movements with an Eye Tribe software, a low-cost eye tracking instrument. Beyond budgetary considerations the instrument was produced by that IT group whose open source GazeTracker software we used previously for our visualization software MapReader [20]. The commercial software and remote tracker works with an average accuracy of less than 0.5 ° of visual angle. The default sampling rate of the instrument (30 Hz) was used, which allowed the further study of fixations.

### C. Procedure

Before the experiment, the participants were assigned randomly into three different experimental groups. These groups differed in the type of exploration and the learning style (active-passive). One group was not given any specific instructions (Free exploration group), the two other groups was given a 5-minute long, guided tour around the city with the only difference of the exploration route, which was indicated in purple on the map of one of them (Guided exploration with route) and not on the other (Guided exploration).

After the exploration the test phase started. Participants had to navigate to find four different shops, approximately at the same distance from each other (knowing the route ∼1.20 minute). Each participant heard four different route descriptions from a matrix of route descriptions, in sum we used nine different combinations of the routes, therefore each route sequence was used at least two times. The route plans showed the optimal way to the shops, considering the shortest distance and the lower complexity (number of turns) of the routes. The directions were given in a verbal form, and if someone forgot it and lost his/her way, they had to go back to the starting point of the section - which was always the end of the previous route – to listen to the directions again. If the participants got lost and was helpless, the experimenter instructed them how to navigate back to the starting point of the actual route, but only after five minutes of ramble.

In the end, after the successful navigation to the last target’s location, the game was over. The participants needed to answer five additional question, about their navigation strategies during the game. We asked them the followings: what type of strategy did they use to navigate in the town; how much did they use the map; how helpful did they find the presence of the map; do they remember the four square and main roads of the map; how would they give a direction to another person from the first and the last shop back to the town center. The whole experiment session lasted approximately 30-40 minutes.

## III. DATA ANALYSIS AND VISUALIZATION

### A. Data preprocessing

At present we could analyze the data collected by the eye tracking device. For the limited accuracy of the instrument mainly fixations were considered in the context of our paradigm. From the heat map displaying accumulated fixations of all participants it seems that our subjects were sporadically looking at the map along the routes, and, if they looked at the cartographic representation their gaze was directed to the central are with central part of the image (with dense street network and symbols). However, during the navigation they mostly looked at the upper part of the screen, the 3D city view, which was intentionally placed at eye-level and slightly above to imitate real world situation, actual map use in the field. These preliminary observations met our a priori expectations.

Next we turned our attention to the shorter periods of switch between the two visual (and conceptual) spaces (Figure 3) and analyzed the data revealing some remarkable aspects of cognitive strategies, namely map reading during spatial movement. These periods are in contrast with the much better studied and understood periods of actual spatial navigation when people do not look at the map but use the internal cognitive apparatus.

For further statistical analysis of the collected eye tracking data we used the R (3.5.0) statistical software, data visualizations are done with ggplot2 by Wickem. Data was merged and loaded into R. First we grouped and categorized fixations to City and Map clusters of fixations using the midline of the screen. We described each cluster with a six dimensional vector; that is

- location of the cluster in terms of City or Map
- duration of cluster
- time of the switch (relative to the start of the task)
- route taken until switch
- route taken during the cluster

Additionally, since gender differences in spatial navigation are well-documented we have taken this as a factor together with the experimental condition group and the ordinality of the current task (number between 1 and 4).

### B. Analysis and Visualization

First, we examined whether the proportion of time spent on viewing the map differs between groups. For this, we modelled the proportion of map watching time (time spent watching the map divided by the total time spent with the task) in function of the group and the gender of the participant. We *did not find difference* between the different experimental conditions (p = 0.7), but we found an effect of *gender* (F (1,51) = 4.75, p =.03): female participants looked at the map significantly shorter overall than male participants. The effect was similarly present both in the exploration (F (1,51) = 4.96, p =.03) and in the test phase (F (1,51) = 3.73, p =.06).

We continued our investigation by looking at the frequency of map watching. This was operationalized by the number of map fixations per second. Here, there was no gender difference (p=.15) but a clear difference between experimental conditions (F (2,51) = 4.22, p =.02): *participants significantly more often looked at the map in the Free exploration* then in the Map-driven exploration with line condition (Tukey HSD diff=0.06, p =.03). The effect was similarly present both in the exploration (F (2,51) = 4.60, p =.01) and in the test phase (F (2,51) = 3.73, p =.03).

As the last step of the statistical analysis, we constructed a general linear model to predict whether a fixation was on the map of on the city. We used logistic link for the modeling, and entered Gender, Experimental group, Task ‘Ordinality’ along with Time of switch, Route taken until the switch, Duration of cluster, Route taken during the cluster as factors in the model. We modelled also the interaction between Experimental Condition and the time and route factors in the model. Since Duration and Route taken during cluster showed a right skewed distribution we entered their log values in the model. The model was only fitted on data from the test phase. The results showed a significant main effect of *Gender* (χ^2^ (1) = 3.70, p=.05); as was earlier shown female participants looked less often at the map. Furthermore, we found and effect of Task ‘Ordinality’ (χ^2^ (1) = 4.13, p=.04). This latter effect was caused by looking significantly more often on the map during later stages of the experiment (Estimate = 0.04, SE = 0.02, z=2.03). Also, a tendency of Route taken until the switch also showed that even within stages people looked more often on the map as time passed (χ^2^ (1) = 2.98, p=.08). The largest effect was Duration of cluster which derives from the clear pattern of watching longer the city than the map (χ^2^ (1) = 82.47, p<.001).

We found also main effect of Experimental group (χ^2^ (1) = 8.17, p=.02), and Route taken during cluster (χ^2^ = (1) = 59.83, p<.001). However, since these factors were also included in a significant interaction (χ^2^ (2) = 8.69, p=.01) we refrain from interpretation of the main effects. The interaction effect was present because participants in the Free exploration group *walked significantly longer distances while looking at the map* as shown by their higher slope both in the Guided exploration (z = 2.487, p =.01) and in the Guided exploration with route (z = 2.547, p =.0) (see Figure 4).

**Figure 4.**
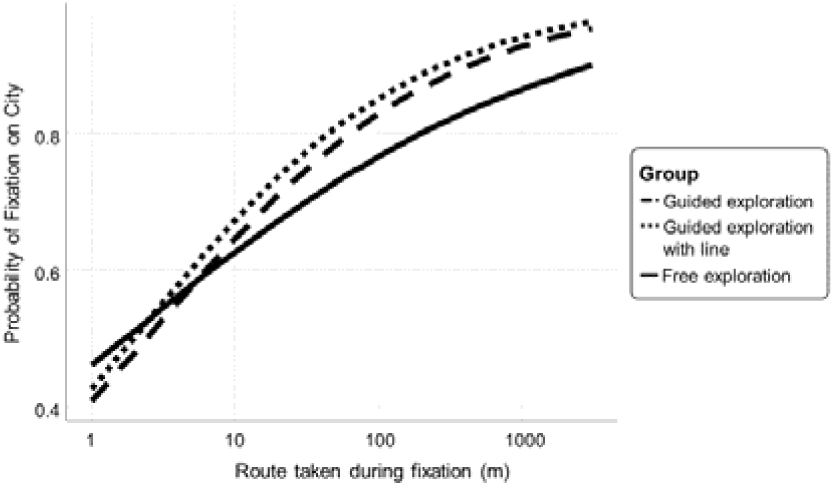
Diagram representing the probability of map use during navigation in the virtual town environment

## IV. CONCLUSIONS

In our study we investigated the role of maps in active and passive spatial knowledge acquisition in large-scale virtual environment. We provided the participants of the experiment a Google style cartographic representation of the city environment, a north-oriented, static map and created a 3D interactive environment where they could navigate. Since our non-immersive virtual environment (VE) did not allow sufficient proprioceptive information the conclusions drawn from the present study have some limitations. Although we were anxious to imitate actual map use situations the experiment could not be realized in real world. However, our ecologically more valid research paradigm, for example the use of a real map with symbolic representation instead of an aerial photo or ground plan made it possible to focus on cognitive features of maps, as the most common visuospatial display in active and passive spatial learning [21].

We concluded to spatial knowledge acquisition based on the proportion of fixations on the map. We found no difference in the performance between the three groups (free exploration, guided exploration, guided with route exploration groups). According to our results, in the route planning task they could navigate to the location of landmarks, and they spent similar amount of time with orientation and wayfinding. The only cases when we found significant increase in visual attention towards and fixation on the map was when guided participants were given a direct map cue in the verbally presented route plan [22]. This indicates that participants indeed tend to use the map to orient themselves in the virtual environment, on the other hand, this intention did not correspond to efficient map usage. The increased frequency of fixation did not lead to better or more accurate wayfinding in test phase since the active learning group did not find the targets’ locations faster or more accurately.

The results of the experiment revealed significant differences in the spatial learning strategies between the guided groups and the free exploration group:

- the more time was spent in the virtual navigation game participants looked more frequently at the map
- although the amount of time spent with map reading was approximately the same at all groups, participants in the free exploration group looked *more frequently* on the map, but they attended it for *shorter periods*
- participants in the free exploration group did not stop when they looked at the map - but kept walking

In conclusion it seems that in time the advantage of the cartographic representation in the acquisition of spatial knowledge and the construction of the cognitive map was realized for all. Although the overall performance was not significantly different in our experiment, participants whose navigation was not influenced by verbal instructions apparently adopted different spatial learning strategies when exploring the virtual world.

Beyond these general observations, however, there is a wide variety of individual spatial abilities, significantly different and alternative learning and navigation strategies, still a huge field that requires future research and analysis [23].

## ACKNOWLEDGMENT

The authors are grateful to the students in the Cognitive Cartography Research Group: Borbála Tölgyesi (cognitive science MSc), Judit Kiss and Veronika Kiss (cartography MSc) and Ádám Bérces (PhD student) for their contribution to the 2016 multidisciplinary research project ‘*The Role of Reference Frame in Visual Cognition*’ at the Department of Cartography and Geoinformatics, supported by Tehetséggondozási Tanács, Eötvös Loránd University.

